# The Gender Attractiveness Gap

**DOI:** 10.1101/2025.05.21.655261

**Authors:** Eugen Wassiliwizky, Brendan P. Zietsch, Fredrik Ullén

**Affiliations:** Department of Cognitive Neuropsychology, Max Planck Institute for Empirical Aesthetics, Frankfurt am Main, Germany; Centre for Psychology and Evolution, School of Psychology, University of Queensland, Queensland, Australia; Department of Neuroscience, Karolinska Institutet, Sweden

**Keywords:** facial attractiveness, gender gap, evolutionary psychology, mate preference, sexual dimorphism

## Abstract

Writers from Darwin to Dawkins have noted that, in humans, women are considered the “beautiful sex,” whereas in most species, it is the males who display more elaborate and visually striking traits. This reversal of typical sex roles is a peculiarity of the human species and has been the focus of extensive theoretical debate. Yet, it has never been systematically examined or empirically verified. Here, we present a comprehensive cross-cultural meta-analysis of same-sex and opposite-sex ratings of facial attractiveness from around the globe. Our findings confirm the existence of a robust “Gender Attractiveness Gap” (GAP), with female faces rated significantly more attractive than male faces across rater genders, cultural backgrounds, and portrayed ethnicities. Notably, the effect is more pronounced among female than male raters, suggesting gender-specific modulation. We show that approximately two-thirds of the GAP is mediated by structural facial sex-typicality. However, this mediation is asymmetric: controlling for facial dimorphism substantially reduces the attractiveness of female—but not male—faces, indicating a specific aesthetic preference for structural femininity by both male and female raters. This suggests that attractiveness judgments are driven by general aesthetic evaluation processes that extend beyond heterosexual mate choice. We also observe a general tendency for greater stringency among male raters. Overall, these findings contribute to evolutionary psychology and social perception, offering new insights into mate selection theories and underscoring the importance of accounting for gender-specific and cultural influences in attractiveness judgments.

## Introduction

The assessment of facial attractiveness is a fundamental aspect of human interaction, often operating subconsciously and continuously in everyday life (Chatterjee et al., 2009; Hahn & Perrett, 2014; Kim et al., 2007; Langlois et al., 2000; Wassiliwizky & Menninghaus, 2021). In particular, the evaluation of facial attractiveness plays a key role in shaping human mate choice (Gangestad & Scheyd, 2005; Little et al., 2011; Rhodes, 2006). A central idea in evolutionary psychology proposes that preferences commonly observed across individuals and populations may have evolved to enhance inclusive fitness—the overall genetic success of an individual through the survival and reproduction of their offspring and kin (Kościński, 2007, 2008; Langlois et al., 2000). Selecting mates with higher genetic quality increases the likelihood of producing healthier and more viable offspring. One prominent example of such a preference is facial averageness, which reflects how closely an individual’s structural facial features align with the population-wide average. Facial averageness is believed to signal genetic quality and resistance to developmental disruptions (Polak, 2003). Thus, by favoring faces with higher averageness, individuals indirectly are assumed to select mates with higher genetic quality, thereby enhancing their reproductive success. As a result, the association of these structural facial features with aesthetic preferences and motivational tendencies observed in contemporary datasets is likely to reflect evolutionary processes shaped over millennia of human history.

Another important line of thought in evolutionary theory highlights that the sexes have different optimal strategies in mate selection, shaped by differences in parental investment (Buss, 1989, 1994; Kokko & Jennions, 2008). In most sexually reproducing species, females have evolved to be the choosy sex, placing selective pressure on males to compete for their attention. Female choice effectively determines which males reproduce and which do not, driving competition among males. This dynamic explains why, in species with sexual dimorphism—where the sexes differ in appearance— males are typically more ornamented and visually striking (Prum, 2017; Ryan, 2018). Darwin’s theory of sexual selection was, in fact, developed to explain such phenomena as the extravagant plumage of male peacocks. Beyond visual traits, these selective pressures also shape elaborate male behaviors, such as singing, courtship dances, and other complex displays intended to captivate potential mates.

However, Darwin himself observed that, in contrast to examples from the animal kingdom, this dynamic appears to be reversed in humans (Darwin, 1871, p.618). In other words, it is women, not men, who are often regarded as the “beautiful sex.” This peculiarity of the human species has been explored by numerous scholars, including Dawkins (1976) and Symons (1979), and has long been considered an evolutionary riddle (Gottschall, 2007). Fixed expressions idealizing female beauty in various languages—such as “the fair sex” in English, “das schöne Geschlecht” in German, “le beau sexe” in French, “il gentil sesso” in Italian, “el sexo bello” in Spanish, “прекрасный пол” in Russian, “美しい性” in Japanese, and “**美丽的性别**” in Chinese— further corroborate this notion.

Surprisingly, despite its long-standing presence in scientific debates and cultural discourse, and an extensive body of empirical work on facial attractiveness over the last decades, the existence of what we can call the Gender Attractiveness Gap (GAP, Gender Attractiveness gaP) has never been systematically verified. Many authors, predominantly men, often assume a shared understanding that women are inherently more attractive than men. However, this assumption raises a critical question: Do both men and women perceive women as more attractive, or does this notion merely reflect a gendered bias among these authors? Without direct empirical evidence, both the true nature and even the very existence of the GAP remain uncertain.

To put the issue on a solid foundation, we conducted a comprehensive meta-analysis of published original datasets of facial attractiveness ratings from the past decade, leveraging the increased availability of shared datasets through open science initiatives. We focused on faces because they have been extensively studied with large samples of stimuli and raters. From a Scopus search using terms related to face and attractiveness, we selected studies that evaluated both male and female images and made their datasets publicly available. We then applied additional filtering, detailed in the Methods section, to include studies that met our predefined criteria. This approach enabled us to compile and harmonize a large, global dataset of male and female raters evaluating same- and opposite-sex facial images. With this harmonized dataset, we rigorously assessed the existence of the GAP and precisely measured its magnitude with robust cross-cultural validity. Moreover, we examined whether structural facial features could account for the GAP.

## Methods

### Literature search and data harmonization

We conducted a Scopus search using the terms ‘attractive’, ‘beauty’, and ‘facial’, employing wildcard operators to capture variations such as ‘attractiveness’, ‘face’, or ‘beautiful’. These terms were required to appear in the title, abstract, or keywords. We selected peer-reviewed studies that (i) used healthy human raters (explicitly identifying as heterosexual in most studies), (ii) evaluated the attractiveness of both male and female faces, and (iii) made their datasets publicly available. From these studies, we included attractiveness rating data for front-view face images with natural or neutral expressions, depicted alone, and without masks or social context. In some cases, this required selecting subsets of data that met these criteria, such as excluding ratings for faces with emotional expressions, medical masks, or other confounding variables.

The final set included contributions from 28 independent studies conducted across more than 50 countries, comprising over 12,500 raters and over 11,000 facial images spanning six ethnic groups (African, Asian, Caucasian, Latino, Middle Eastern, and Multi-Ethnic). Almost all included studies used ecologically valid real face images, with one exception where computer-generated faces were used. In all included studies, both the gender of the raters and that of the portrayed individuals were conceptualized in binary terms as male and female in the original data. Consequently, our findings can only be generalized to individuals who identify as binary. None of the selected studies explicitly examined gender-based differences. Rather, they addressed research questions that required collecting facial attractiveness ratings, such as the effects of emotional expressions (Ebner et al., 2018), sex-typicality (Fiala et al., 2021), or facemasks (Hewer & Lewis, 2024) on perceived attractiveness. Table 1 provides an overview of all studies included in the harmonized dataset, detailing raters, stimuli, additional measures, and the rating conditions implemented in their experimental designs. Since our statistical model operated on the full dataset, we also included studies that did not employ a fully crossed design—specifically, four datasets containing cross-gender ratings (Studies 4, 10, 15, and 27 in Table 1).

**Table 1.** List of studies incorporated into the harmonized dataset, including the (selected) number of raters and stimuli. Female images are represented by ‘F’, and male images by ‘M’. Female raters are represented by ‘f’, and male raters by ‘m’. The middle dot (•) indicates that male and female ratings have been averaged in the published dataset. The last column lists physical facial properties provided by the authors of some studies, including morphometric features (AVG = facial averageness, SShD = sexual shape dimorphism), as well as the depicted person’s age. Measured characteristics not relevant to the current analysis, such as body mass index, skin luminance, or eye color, are not listed.

| Harmonized Dataset |  |  |  |  |  |  |
| --- | --- | --- | --- | --- | --- | --- |
| No. | Study | N<sub>Stim</sub> | N<sub>Rater</sub> | Data Type | Conditions | Facial Features |
| 1 | Bowdring et al. 2021 | 216<br>(108 F, 108 M) | 181<br>(106 f, 75 m) | raw | F:f, M:f,<br>F:m, M:m |  |
| 2 | Chen et al. 2021 | 110<br>(90 F, 20 M) | 2082<br>(1133 f, 949 m) | aggr | F:•,<br>M:• |  |
| 3 | Ebner et al. 2018 | 342<br>(170 F, 172 M) | 154<br>(76 f, 76 m) | aggr | F:f, M:f,<br>F:m, M:m |  |
| 4 | Fiala et al. 2021 | 709<br>(352 F, 357 M) | 2455<br>(999 f, 1456 m) | raw* | M:f,<br>F:m | AVG, SShD,<br>Age |
| 5 | Germine et al. 2015 | 260<br>(132 f, 128 m) | 1548<br>(1174 f, 374 m) | raw | F:f, M:f,<br>F:m, M:m |  |
| 6 | Gouda-Vossos et al. 2016 | 32<br>(15 F, 17 M) | 940<br>(668 f, 272 m) | raw | F:f, M:f,<br>F:m, M:m | Age |
| 7 | Hewer et al. 2024 | 48<br>(24 F, 24 M) | 85<br>(76 f, 9 m) | raw | F:f, M:f,<br>F:m, M:m |  |
| 8 | Jaeger et al. 2019 | 1020<br>(529 F, 491 M) | 1336<br>(660 f, 676 m) | raw* | F:f, M:f,<br>F:m, M:m | Age |
| 9 | Kleisner et al. 2024 | 167<br>(87 F, 80 M) | 777<br>(577 f, 200 m) | raw* | F:f, M:f,<br>F:m, M:m | AVG, SShD,<br>Age |
| 10 | Kočnar et al. 2019 | 120<br>(60 F, 60 M) | 762<br>(447 f, 315 m) | raw* | M:f,<br>F:m | AVG, SShD,<br>Age |
| 11 | Leger et al. 2024 | 100<br>(50 F, 50 M) | 400<br>(200 f, 200 m) | raw | F:f, M:f,<br>F:m, M:m | AVG, SShD,<br>Age |
| 12 | Lindeberg et al. 2019 | 167<br>(84 F, 83 M) | 43<br>(29 f, 14 m) | raw | F:f, M:f,<br>F:m, M:m |  |
| 13 | Mitchem et al. 2015 | 521<br>(276 F, 245 M) | 23<br>(13 f, 10 m) | raw | F:f, M:f,<br>F:m, M:m | Age |
| 14 | Nakamura et al. 2019 | 140<br>(70 F, 70 M) | 48<br>(24 f, 24 m) | aggr | F:•,<br>M:• |  |
| 15 | Pavlovič et al. 2023 | 93<br>(33 F, 60 M) | 509<br>(315 f, 194 m) | raw | M:f,<br>F:m | AVG, SShD,<br>Age |
| 16 | Qi et al. 2024<br>(Studies 1 + 2) | 119<br>(61 F, 58 M) | 140<br>(74 f, 66 m) | raw | F:f, M:f,<br>F:m, M:m |  |
| 17 | Sano et al. 2023<br>(Study 1a) | 5500<br>(2750 F, 2750 M) | 60<br>(30 f, 30 m) | aggr | F:•,<br>M:• |  |
| 18 | Sano et al. 2023<br>(Study 1b) | 300<br>(150 F, 150 M) | 63<br>(37 f, 26 m) | raw | F:f, M:f,<br>F:m, M:m |  |
| 19 | Sutherland et al.<br>2017 | 64<br>(32 F, 32 M) | 16<br>(8 f, 8m) | aggr | F:•,<br>M:• |  |
| 20 | Sutherland, Rhodes<br>et al. 2020 | 200<br>(100 F, 100 M) | 24<br>(12 f, 12 m) | raw | F:f, M:f,<br>F:m, M:m |  |
| 21 | Sutherland, Burton<br>et al. 2020 (lab) | 34<br>(10 F, 24 M) | 214<br>(151 f, 63 m) | raw | F:f, M:f,<br>F:m, M:m |  |
| 22 | Talamas et al.<br>2016 | 107<br>(69 F, 38 M) | 32<br>(11 f, 21 m) | raw* | F:f, M:f,<br>F:m, M:m | Age |
| 23 | Torrance et al.<br>2014 | 100<br>(50 F, 50 M) | 303<br>(215 f, 88 m) | aggr | F:•,<br>M:• | Age |
| 24 | Wassiliwizky et al.<br>2023 | 380<br>(186 F, 194 M) | 80<br>(42 f, 38 m) | raw | F:f, M:f,<br>F:m, M:m | Age |
| 25 | Wassiliwizky et al.<br>(forthcoming) | 126<br>(63 F, 63 M) | 112<br>(57 f, 55m) | raw | F:f, M:f,<br>F:m, M:m |  |
| 26 | Wassiliwizky et al.<br>(forthcoming) | 16<br>(8 F, 8 M) | 84<br>(54 f, 30 m) | raw | F:f, M:f,<br>F:m, M:m |  |
| 27 | Zhang et al. 2019 | 200<br>(100 F, 100 M) | 200<br>(100 f, 100 m) | raw | M:f,<br>F:m |  |
| Sum |  | **11,191**<br>**(5659 F, 5532 M)** | **12,671**<br>**(7288 f, 5381 m)** |  |  |  |

\*for this study, we obtained the raw data from the corresponding authors; the published dataset is aggregated per stimulus

We z-standardized the ratings before incorporating them into the harmonized dataset and computed gender-specific stimulus means, as not all studies provided the raw data from individual raters. Re-analysis of studies with raw data are provided in the Supplemental Material (Table S1, Figure S1). They confirmed identical result patterns, showing that aggregation did not affect the results. Data of five studies where ratings had already been averaged into gender-specific stimulus means were subset and analyzed separately (hereafter referred to as the Subset with Collapsed Ratings).

Figure 1 presents the distribution and normality assessment of attractiveness scores, aggregated at the stimulus level, for both female (N = 5,659) and male (N = 5,532) images of the entire harmonized dataset. Notably, even at a descriptive level, the male distribution appears shifted leftward, suggesting the presence of the GAP.

**Fig. 1.**
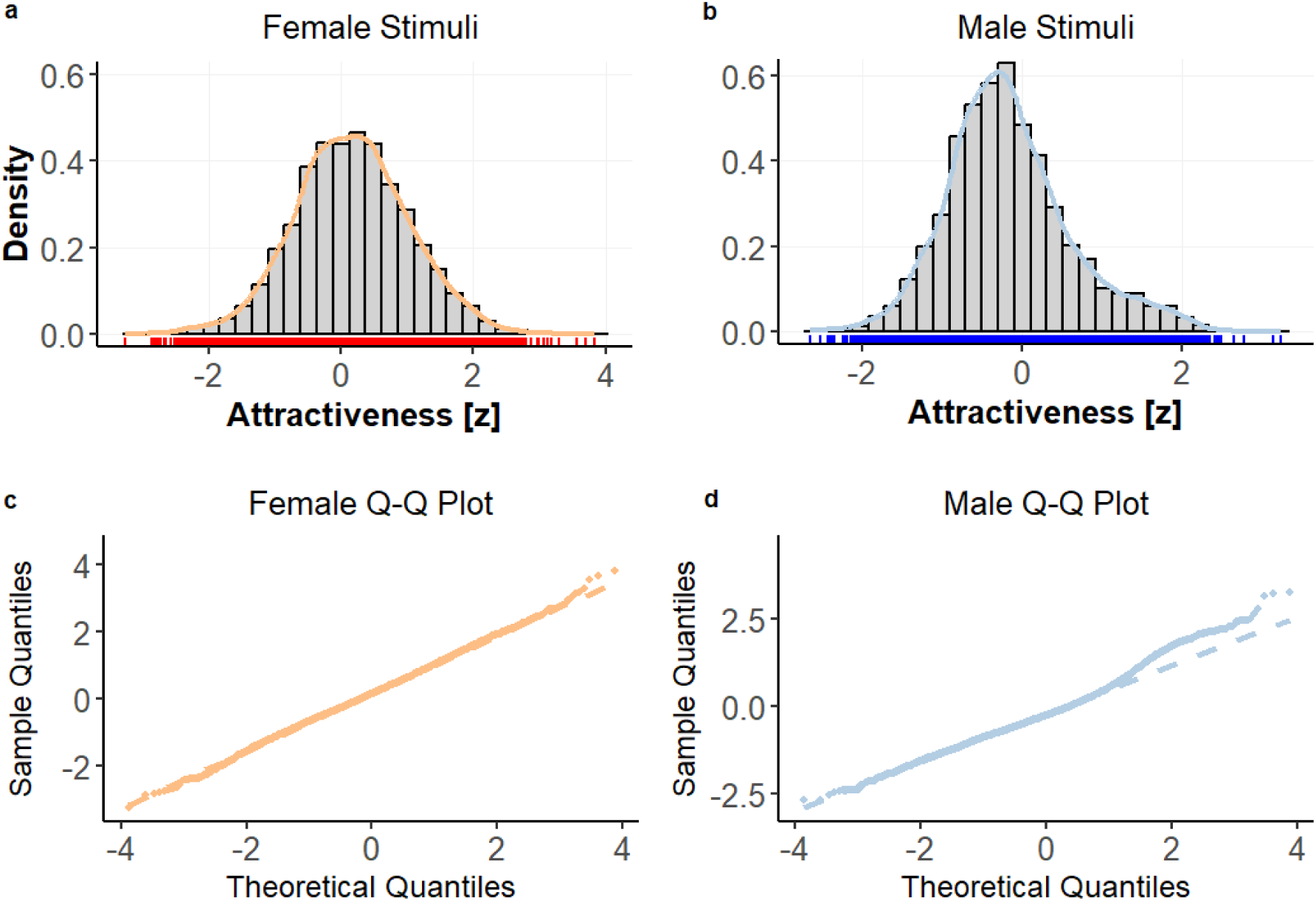
Distribution and normality assessment of attractiveness scores in the harmonized dataset. The upper row displays histograms with overlaid density plots for female (a) and male (b) stimuli. Rug plots (red and blue ticks along the x-axis) represent data points for individual stimuli. The lower row presents Q-Q plots assessing normality for female (c) and male (d) stimuli, with sample quantiles plotted against theoretical normal quantiles. Dashed lines indicate expected normal distributions. While both distributions align closely with normality in the central range, male stimuli exhibit a slight right-tail deviation.

### Statistical Analysis

#### Main Analysis

We employed linear multi-level modeling (LMM) to analyze how attractiveness ratings (Level 1) varied based on the gender of the portrayed individual (StimGen), the rater’s gender (RaterGen), and their interaction. Facial stimuli were treated as Level 2 units in the hierarchical model, with random intercepts included to account for baseline differences in attractiveness across images. The following model was tested: Attractiveness_i_ = β_0_ + β_1_ StimGen_ij_ + β_2_ RaterGen_ij_ + β_3_ (StimGen_i_ x RaterGen_i_) + ε_i_. For studies where ratings were collapsed across RaterGen (i.e., five studies indicated by a middle dot in Table 1), attractiveness was predicted solely by the image’s gender: Attractiveness_ij_ = β_0_ + β_1_ StimGen_i_ + ε_i_.

#### Ubiquity Analyses

After the main analysis, we tested the consistency of the GAP across both the cultural backgrounds of the raters and the ethnicities of portrayed individuals. To this end, we conducted two analyses. In the first LMM, we examined whether the effect of StimGen on attractiveness varied across raters’ cultural backgrounds by modeling their interaction: Attractiveness_i_ = β_0_ + β_1_ (StimGen_i_ x CultBack_i_) + ε_i_. Given that our harmonized dataset had participants from over 50 countries, we grouped them into ten broader cultural regions as follows:

− Anglophone Countries (Australia, Canada, New Zealand, UK, USA)
− Asia Eastern (China, Japan, South Korea, Taiwan, Vietnam)
− Asia Southern (India, Malaysia, Singapore)
− Europe Eastern (Croatia, Czech Republic, Estonia, Hungary, Moldova, Poland, Romania, Russia, Serbia, Slovakia, Ukraine)
− Europe Northern (Denmark, Finland, Norway, Sweden)
− Europe Southern (Greece, Italy, Portugal, Spain)
− Europe Western (Austria, Belgium, France, Germany, Ireland, Netherlands, Switzerland)
− Latin America (Argentina, Brazil, Chile, Colombia, Mexico)
− Middle East (Egypt, Emirates, Iran, Israel, Turkey)
− Sub-Saharan Africa (Cameroon, Namibia, South Africa)

In the second model, we examined whether the effect of StimGen on attractiveness varied based on the portrayed individual’s ethnicity (categorized as African, Asian, Caucasian, Latino, Middle Eastern, or Multi-Ethnic) by modeling their interaction: Attractiveness_i_ = β_0_ + β_1_ (StimGen_i_ x StimEthnicity_i_) + ε_i_. For both models, stimuli were again treated as Level 2 units. In pairwise post-hoc analyses, we applied a Bonferroni correction to control for alpha inflation. Ethnic categorization of portrayed individuals was obtained from the original data, while raters’ cultural backgrounds were determined by their country of residence. Stimuli for which the ethnic membership of portrayed individuals could not be reconstructed were excluded from the second analysis (i.e., Studies 1, 6, 13, 20, and 21 in Table 1).

#### Morphometric Mediation Analysis

In an additional series of analyses, we examined the extent to which the GAP was mediated by structural features that define male and female faces. Facial morphometry provides a valuable framework for investigating this question, as it quantifies objective features, thereby helping to explain subjective perceptions. In our harmonized dataset, multiple studies contributed various facial metrics alongside ratings of perceived attractiveness. The key metric for our case is sexual shape dimorphism, which is a holistic measure of sex-typicality. It is calculated for each face using statistical shape analysis of facial landmarks, assessing their Procrustes fit with population-based male and female configurations (Holzleitner et al., 2014; Lee et al., 2014). The resulting score places a face along a female-male continuum, with higher positive values indicating a more masculine facial shape, higher negative values indicating a more feminine shape, and scores around zero reflecting androgyny. In general, sex-typicality is preferred by the opposite gender—that is, greater masculinity in male faces and greater femininity in female faces are typically associated with higher attractiveness (Bronstad et al., 2008; Muñoz-Reyes et al., 2015; Penton-Voak et al., 2001; Perrett et al., 1998; Rhodes et al., 2000). However, the relationship varies across cultural and socioeconomic contexts, particularly with regard to female preference for masculinity in male faces (Fiala et al., 2021; DeBruine et al., 2010; Kleisner et al., 2024; Marcinkowska et al., 2019; Rennels et al., 2008).

With sexual shape dimorphism being linked to both the gender of the image and perceived attractiveness, we were able to perform a mediation analysis to determine how much of the GAP is explained by structural sex-typicality. To investigate this mediating relationship as purely as possible, we first partialed out two other well-known influences on attractiveness—facial maturity and facial averageness. We refer to the resulting measure as residualized attractiveness. Facial maturity, closely linked to fertility and health, was estimated by the depicted individual’s age and is typically negatively related to attractiveness across individuals and populations (Buss, 1989; Deffenbacher et al., 1998; Ebner et al., 2018; Maestripieri et al., 2014). Facial averageness was computed in the original studies similarly to sexual shape dimorphism by projecting individual facial landmarks onto a population-based average configuration and quantifying the resulting Procrustes residuals (Lee et al., 2016). This method captures the extent to which individual facial landmarks deviate from the average face, with larger scores indicating a less average (or more distinct) face, which typically receives lower attractiveness ratings (Apicella et al., 2007; Fiala et al., 2021; Kleisner et al., 2024; Komori et al., 2009; Lee et al., 2016; Rhodes et al., 2001). In our dataset, too, we observed that attractiveness ratings were influenced by all three morphometric measures: deviation from averageness (β_1j_ *= −0.145 [−0.156, −0.133]*, *t = −*24.033, *p < 0.001*, *d = −0.18*), facial maturity (β_2j_ *= −0.113 [−0.124, −0.102]*, *t = −*20.528, *p < 0.001*, *d = −0.14*), and sexual shape dimorphism (β_3j_ *= −0.15 [−0.166, −0.132]*, *t = −*17.411, *p < 0.001*, *d = −0.19*). Model fit improved incrementally with the inclusion of each predictor, with the best performance observed when all three factors were included, highlighting their combined relevance in explaining attractiveness ratings (see Table S3).

For all morphometric analyses, we utilized a reduced dataset comprising the necessary metrics (Table 1), which we z-standardized for the analyses. Since we had access to the raw data from all respective studies, we used these original raw data for the morphometric analyses, incorporating random effects for both images and raters to obtain the most accurate parameter estimates while accounting for inter-individual differences among raters. Replication of the mediation analysis using aggregated data produced similar results but with wider confidence intervals due to the naturally smaller sample size (Figure S2).

All statistical analyses were conducted in R, version 4.4.1 (R Core Team, 2024), using the *lme4* package, version 1.1.35.5 (Bates et al., 2015). Parameter estimates were obtained using the restricted maximum likelihood (REML) estimation method. P-values were calculated using the *lmerTest* package (Kuznetsova et al., 2017). Following recent recommendations (Luo et al., 2021), we also report profile likelihood confidence intervals with 95% boundaries (in square brackets following each parameter estimate). These intervals were computed using the *confint* function within the lme4 package. Effect sizes are reported as d measures, following Westfall et al.’s recommendation, and were calculated as the estimate for the fixed effect divided by the square root of the pooled variance of random effects (Westfall et al., 2014). Westfall’s d effect sizes can be interpreted using the same conventions as Cohen’s d, with the typical benchmarks being: small effect (d ≈ 0.2), medium effect (d ≈ 0.5), and large effect (d ≈ 0.8). For visualizing the results, we calculated estimated marginal means (EMMs) with 95% confidence intervals as error bars (Searle et al., 1980), utilizing the *emmeans* package, version 1.10.3 (Lenth et al., 2023).

All methods and procedures were approved by the Ethics Committee of the Max Planck Society (approval number 2017_12). All scripts and data required to reproduce the results, figures, and tables are available on the Open Science Framework [link follows].

## Results

### Main Analysis

Modeling the harmonized dataset revealed three significant findings (Table 2 left; Figure 2a). First, there was a main effect of StimGen, indicating that male faces generally received lower attractiveness ratings than female faces (β_1_ *= −0.414 [−0.451, −0.377]*, *t = −21.672*, *p < 0.001*, *d = −0.59*). This lends empirical support to the concept of a GAP, first noted by Darwin and later expanded upon by other evolutionary theorists. Westfall’s *d* of −0.59 indicated that this effect is of medium to large effect size. The GAP was also evident in the subset of five studies (Table 2 right, Figure 2b), where ratings were aggregated across the raters’ gender (β_1_ *= −0.276 [−0.326, −0.226], t = −10.831*, *p < 0.001*, *d = −0.28*). Notably, during the collection of datasets for our meta-analysis, we found that the GAP was present in nearly all individual study datasets, with only two exceptions: one study where attractiveness ratings were heavily skewed toward the low end of the scale for both portrayed genders (Study 21, Table 1) and another that used computer-generated faces with an oval-shaped head structure (Study 14, Table 1).

**Fig. 2.**
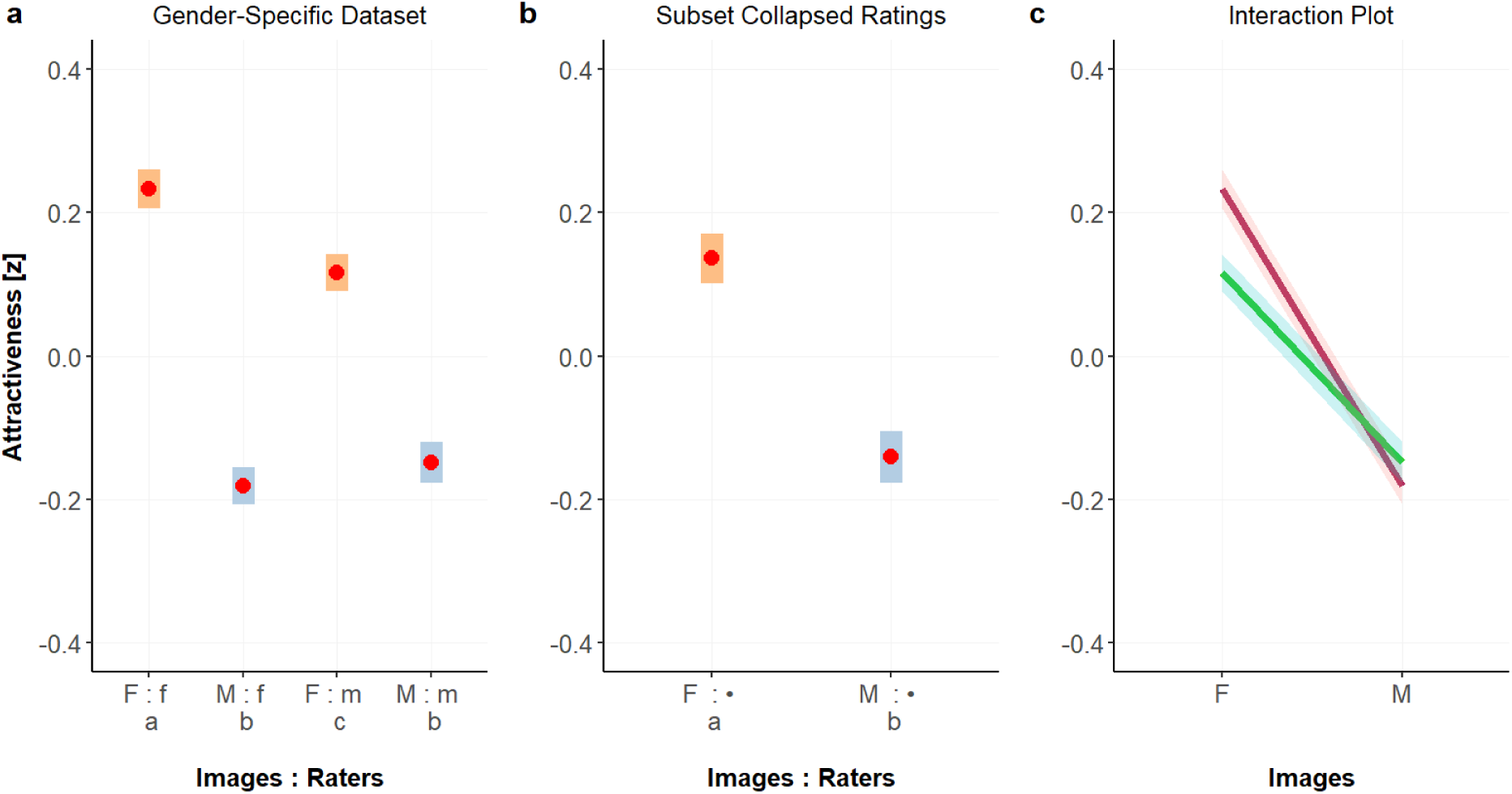
Meta-analysis results for the LMM of the harmonized dataset. Bars depict estimated marginal means (EMMs), including their 95% confidence intervals. Panel **a** shows the results for the Gender-Specific Dataset, where ratings were provided separately by male and female raters. The gender of the facial image is indicated in capital letters, while the gender of the rater is shown in lowercase letters, e.g., in condition ‘F : m’, female faces were rated by male raters. Note that: (1) male faces (in blue) generally receive lower attractiveness ratings (Gender Attractiveness Gap), (2) male raters tend to assign lower ratings overall (Gender Rating Gap), and (3) the discrepancy between female and male faces is more pronounced among female raters. Within each panel, conditions marked with different subscript letters are significantly different at a p-value of < 0.001. For statistical details, please refer to Table 2. Panel **b** displays the EMMs for the Collapsed Ratings Dataset, a subset of five studies where ratings from male and female raters were averaged per image, indicated by the middle dot. Panel **c** depicts the interaction effect between StimGen (on the x-axis) and RaterGen (line colors: maroon for female raters, green for male raters) for the Gender-Specific Dataset. Lines represent predicted attractiveness scores, the shaded ribbons indicate 95% confidence intervals.

**Table 2.** LMM results for the harmonized dataset. In the left part, attractiveness was predicted based on both the image’s gender (StimGen: M, F) interacting with the rater’s gender (RaterGen: m, f). In the right part, LMM results for the subset where ratings were collapsed across rater gender are presented. In this model, attractiveness was predicted only by the image’s gender. For both analyses, stimuli were treated as Level 2 units.

| *Gender-Specific Dataset* |  |  |  | *Subset with Collapsed Ratings* |  |  |
| --- | --- | --- | --- | --- | --- | --- |
| *Predictors* | *Estimates*<br>*[95%-CI]* | *t* | *p* | *Estimates*<br>*[95%-CI]* | *t* | *p* |
| **Fixed Effects** |  |  |  |  |  |  |
| $\beta_0$<br>(Intercept) | 0.233<br>[0.206, 0.26] | 16.749 | <0.001 | 0.135<br>[ 0.1, 0.171 ] | 7.559 | <0.001 |
| $\beta_1$<br>StimGen:M | -0.414<br>[-0.451, -0.377] | -21.672 | <0.001 | -0.28<br>[-0.326, -0.226] | -10.831 | <0.001 |
| $\beta_2$<br>RaterGen:m | -0.117<br>[-0.143, -0.092] | -8.996 | <0.001 | | | |
| $\beta_3$<br>StimGen:M x<br>RaterGen:m | 0.149<br>[0.112, 0.186] | 7.915 | <0.001 | | | |
| **Random Effects** |  |  |  |  |  |  |
| $\epsilon_i$ | | 0.255 | | | 0.546 | |
| $V_0$ | | 0.239<sub>Stim</sub> | | | 0.435<sub>Stim</sub> | |
| ICC |  | 0.484 |  |  | 0.443 |  |
| N |  | 5396<sub>Stim</sub> |  |  | 5863<sub>Stim</sub> |  |
| Observations |  | 13,354 |  |  | 6022 |  |
| Conditional R<sup>2</sup> /<br>Marginal R<sup>2</sup> |  | 0.513 / 0.057 |  |  | 0.454 / 0.019 |  |

Second, we observed a small effect for RaterGen, with male raters generally giving lower attractiveness ratings than female raters (β_2_ *= −0.117 [−0.143, −0.092], t = −8.996, p < 0.001*, *d = −0.16*). Depending on the reference group, this bias can be described as either a male stringency bias or a female benevolence bias. A related phenomenon, referred to as the Gender Rating Gap (GRG), was recently documented in a large-scale analysis of online reviews across various online platforms (Bayerl et al., 2024). Specifically this study found a systematic pattern in which women consistently provided more favorable ratings than men.

Third, there was a significant interaction between the image’s and the rater’s gender, indicating that the discrepancy in ratings for female and male faces depended on the rater’s own gender (β_3_ *= 0.149 [0.112, 0.186]*, *t = 7.915*, *p < 0.001*, *d = 0.21,* representing a small effect size). Female raters exhibited a larger difference in their ratings of male and female faces compared to male raters, leading to a more pronounced GAP for the female raters. As illustrated in Figure 2a, this is reflected in the vertical difference between ratings of male and female faces being wider for female raters (left two bars, conditions ‘F : f’ and ‘M : f’) than for male raters (right two bars, conditions ‘F : m’ and ‘M : m’).

The interaction plot in Figure 2c further illustrates how the image’s and the rater’s genders jointly influence attractiveness ratings. Both lines, representing the ratings by female raters (maroon) and male raters (green), slope downward from left to right, reflecting the overall GAP—male faces were rated lower across the board compared to female faces. The steeper slope of the maroon line highlights that female raters differentiate more strongly between male and female faces compared to male raters.

### Ubiquity of the GAP

The results of the ubiquity analyses revealed significant interactions of the GAP with both the cultural background of raters (*F_19,11753_ = 30.902, p < 0.001, η_p_² = 0.029, f^2^ = 0.03*, representing a small effect size) and portrayed ethnicity (*F_11,10207_ = 42.538, p < 0.001, η_p_² = 0.028, f^2^ = 0.029*, also representing a small effect size). Figure 3 illustrates how the GAP varies across these two factors. For instance, in the Latin American context, both raters and Latino individuals exhibited one of the strongest GAP effects. In contrast, in the African context, no significant GAP was observed for either raters or portrayed individuals, and in both cases, the trend even appeared to reverse direction. However, more data from the African continent is needed to draw reliable conclusions. Additionally, the original studies revealed a strong overlap between raters’ cultural backgrounds and the faces they evaluated. That is, in Western countries, participants predominantly rated Caucasian faces, while in Asian countries, most portrayed individuals were of Asian origin, followed by Caucasians. Across all cultural regions, Caucasian faces were the only widely represented ethnicity (see Table S2 for a cross-tabulation). To address this sampling bias, future studies could aim for broader ethnic representation within cultural regions.

**Fig. 3.**
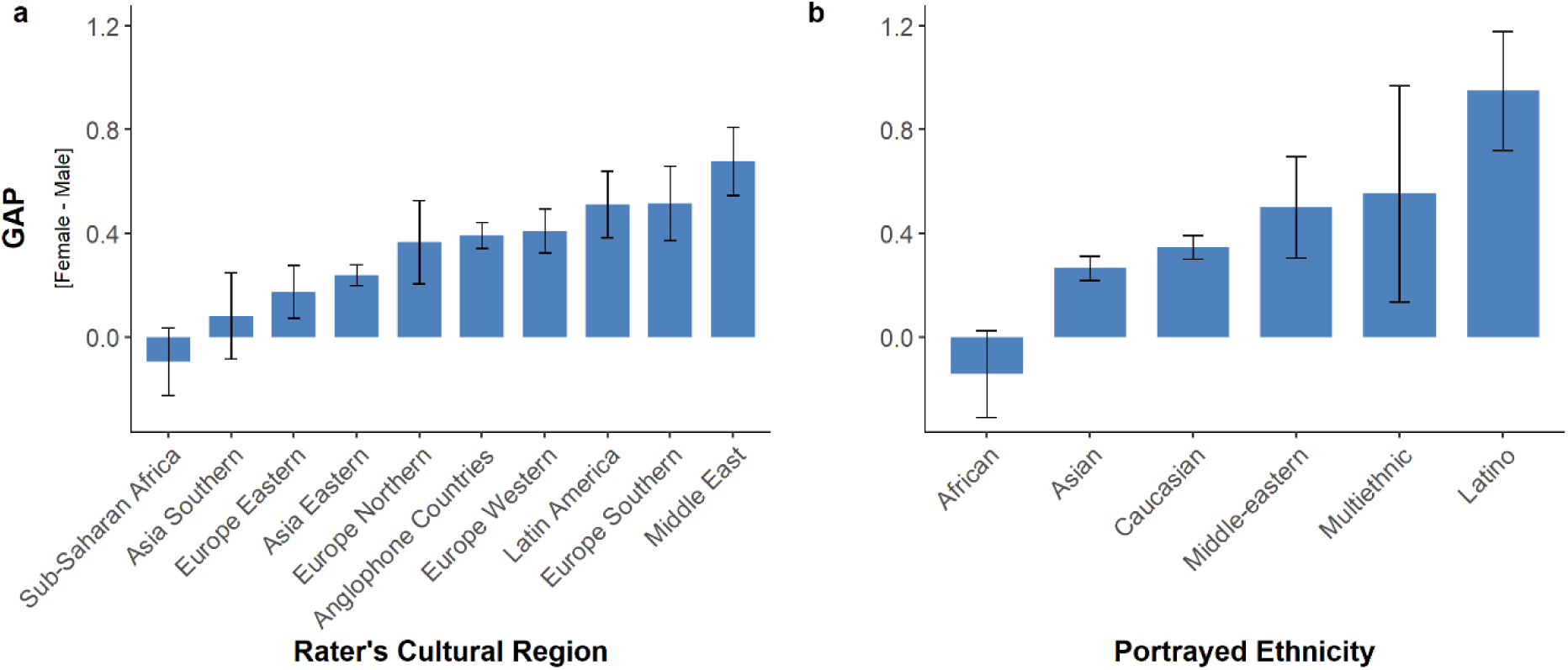
Ubiquity of the GAP. Panel **a** shows the differences in the attractiveness EMMs (z-standardized) across the cultural backgrounds of the raters, ordered by the size of the difference. The error bars reflect 95% confidence intervals. Positive values indicate that female images received higher ratings than male images. The GAP was significant (at p < 0.001) in most regions, except for Sub-Saharan Africa and Southern Asia, where no significant differences were observed. Panel **b** shows the differences in attractiveness ratings across portrayed ethnicities, also ordered by the size of the difference. The GAP remained consistent across all ethnic groups (p < 0.001), except for African faces, where no significant difference was observed. Wider confidence intervals generally correspond to smaller representation within the dataset (cf. Table S2).

### Morphometric Mediation Analyses

The multilevel mediation analysis investigated whether the relationship between StimGen and residualized attractiveness ratings was mediated by the structural sex-typicality of faces, while at the same time accounting for the moderating effect of RaterGen. Detailed results are presented in Table 3 and illustrated in Figure 4. The total effect model (Table 3 path c, Figure 4c) revealed the impact of StimGen (β_1j_ *= −0.372 [−0.441, −0.302]*, *t = −*10.544, *p < 0.001*, *d = −0.47*) and RaterGen (β_2j_ *= −0.186 [−0.244, −0.128]*, *t = −*6.312, *p <* 0.001, *d = −0.24*) on residualized attractiveness, as well as their interaction (β_3j_ *= 0.199 [0.178, 0.219]*, *t =* 18.697, *p <* 0.001, *d = 0.25*). This outcome pattern, including effect sizes, is consistent with our main analysis performed on the entire harmonized dataset (Table 2, Figure 2a).

**Fig. 4.**
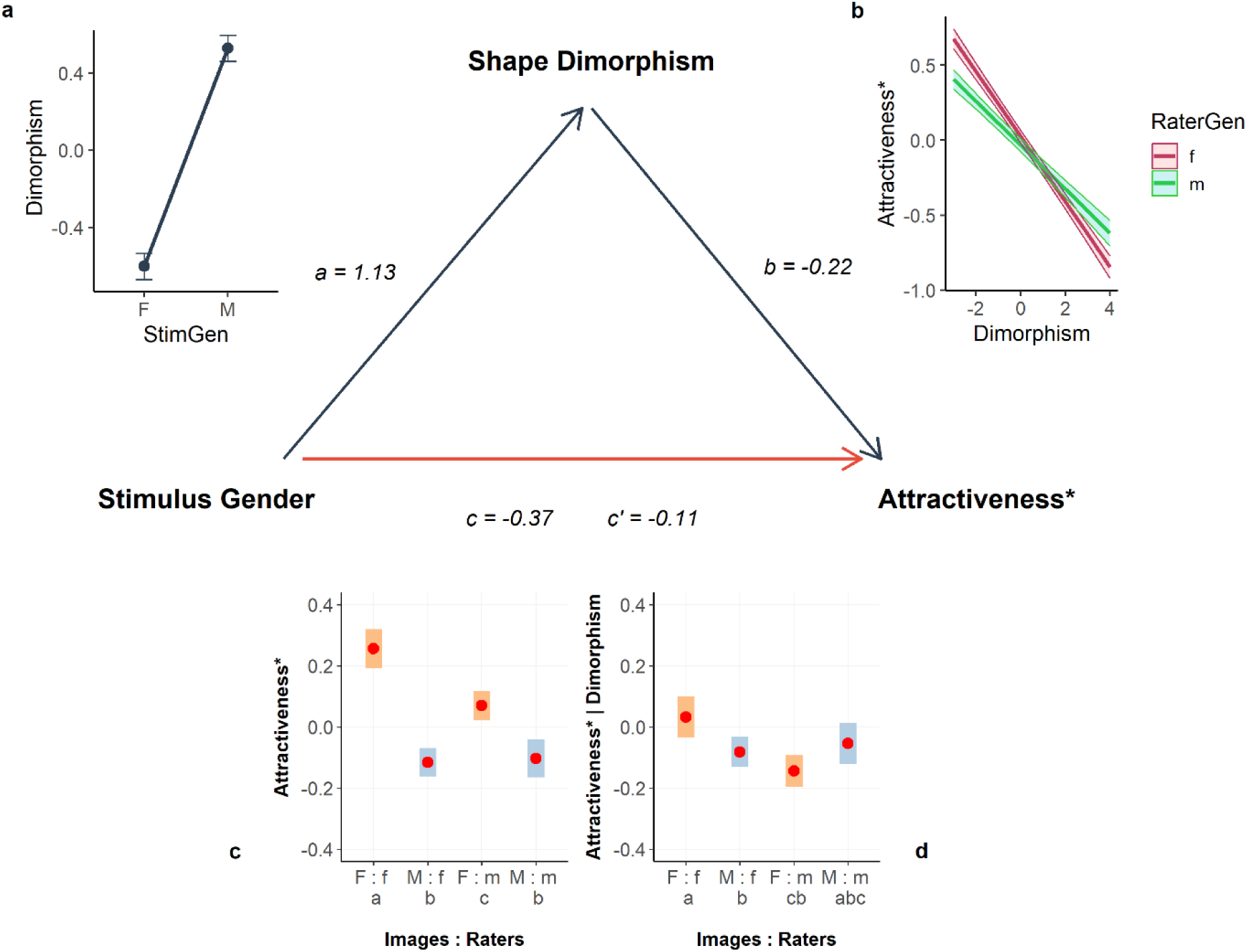
Mediation path model and estimated effect plots. This multilevel mediation analysis examined how the effect of StimGen on attractiveness is mediated by sexual shape dimorphism, while controlling for RaterGen. The influence of averageness and youthfulness has been regressed out from attractiveness, as indicated by the asterisk (*). Path coefficients (a, b, c, c’) are displayed along the respective arrows. Estimated effects plots (a–d) illustrate the modeled relationships between variables: (**a**) Effect of StimGen on dimorphism, with higher values indicating more masculine facial features (a path). (**b**) Effect of dimorphism on residualized attractiveness (b path), separated for female and male raters. (**c**) Total effect of StimGen on residualized attractiveness (c path) showing the GAP, the GRG, and the interaction effect. (**d**) Direct effect of StimGen on residualized attractiveness, after controlling for dimorphism (c’ path). Bars represent EMMs, including their 95% confidence intervals. Within each plot, conditions marked with different subscript letters are significantly different at a p-value of < 0.001. The weakened path strength indicates a partial mediation, with the mediator’s impact observed primarily for female images (orange bars).

**Table 3.** Multi-level mediation analysis. Results on the relationship between StimGen and attractiveness (residualized for averageness and youthfulness), mediated by sexual shape dimorphism. Random effects include within-stimulus and within-rater variability. For a visualization, see Figure 4.

| *Predictors* | *path a* |  |  | *path b* |  |  | *path c* |  |  | *path c'* |  |  |
| --- | --- | --- | --- | --- | --- | --- | --- | --- | --- | --- | --- | --- |
|  | *Estimates*<br>[95%-CI] | *t* | *p* | *Estimates*<br>[95%-CI] | *t* | *p* | *Estimates*<br>[95%-CI] | *t* | *p* | *Estimates*<br>[95%-CI] | *t* | *p* |
| **Fixed Effects** |  |  |  |  |  |  |  |  |  |  |  |  |
| $\gamma_{00}$<br>(Intercept) | -0.602<br>[-0.669, -0.534] | -17.41 | <0.001 | 0.024<br>[-0.018, 0.065] | 1.112 | 0.266 | 0.256<br>[0.192, 0.32] | 7.862 | <0.001 | 0.031<br>[-0.034, 0.098] | 0.930 | 0.352 |
| $\beta_{ij}$<br>StimGen:M | 1.13<br>[1.036, 1.225] | 23.42 | <0.001 | | | | -0.372<br>[-0.441, -0.302] | -10.544 | <0.001 | -0.113<br>[-0.187, -0.039] | -3.021 | 0.002 |
| $\beta_{ij}$<br>RaterGen:m | | | | -0.058<br>[-0.109, -0.006] | -2.189 | 0.028 | -0.186<br>[-0.244, -0.128] | -6.312 | <0.001 | -0.177<br>[-0.236, -0.117] | -5.816 | <0.001 |
| $\beta_{ij}$<br>Dimorphism | | | | -0.217<br>[-0.233, -0.201] | -26.175 | <0.001 | | | | -0.298<br>[-0.322, -0.275] | -25.067 | <0.001 |
| $\beta_{ij}$<br>StimGen:M x<br>RaterGen:m | | | | | | | 0.199<br>[0.178, 0.219] | 18.697 | <0.001 | 0.204<br>[0.172, 0.237] | 12.377 | <0.001 |
| $\beta_{ij}$<br>StimGen:M x<br>Dimorphism | | | | | | | | | | 0.240<br>[0.205, 0.275] | 13.613 | <0.001 |
| $\beta_{ij}$<br>RaterGen:m x<br>Dimorphism | | | | 0.071<br>[0.061, 0.081] | 13.303 | <0.001 | | | | 0.001<br>[-0.018, 0.021] | 0.147 | 0.883 |
| $\beta_{ij}$<br>StimGen:M x<br>RaterGen:m x<br>Dimorphism | | | | | | | | | | -0.009<br>[-0.041, 0.021] | -0.612 | 0.54 |

| Random Effects |  |  |  |  |
| --- | --- | --- | --- | --- |
| $\varepsilon_{ij}$ | 0.026 | 0.45 | 0.451 | 0.448 |
| $V_{i0}$ | 0.633<sub>Stim</sub> | 0.181<sub>Stim</sub> | 0.176<sub>Stim</sub> | 0.181<sub>Stim</sub> |
| $V_{0j}$ | | 0.365<sub>Rater</sub> | 0.366<sub>Rater</sub> | 0.366<sub>Rater</sub> |
| ICC | 0.96 | 0.549 | 0.546 | 0.549 |
| N | 1087<sub>Stim</sub> | 1087<sub>Stim</sub> / 4420<sub>Rater</sub> | 1087<sub>Stim</sub> / 4420<sub>Rater</sub> | 1087<sub>Stim</sub> / 4420<sub>Rater</sub> |
| Observations | 188,696 | 186,779 | 186,779 | 186,779 |
| Conditional R<sup>2</sup> /<br>Marginal R<sup>2</sup> | 0.973 / 0.324 | 0.564 / 0.034 | 0.555 / 0.019 | 0.567 / 0.039 |

The effect of the independent variable StimGen on the mediator Sexual Shape Dimorphism (Table 3 path a, Figure 4a) showed a strong association between StimGen and structural sex-typicality of faces (β_1j_ *= 1.13 [1.036, 1.225]*, *t = 23.42*, *p < 0.001*, *d = 1.39)*. The positive association between StimGen (women coded as 0, men coded as 1) and the objective measure of sexual shape dimorphism (higher values indicate structural masculinity) confirmed that the morphometric measure effectively captured male and female facial features. The large Westfall’s d effect size measure further supported this conclusion. RaterGen and their random effects were not modeled in this path, as it concerns exclusively stimulus features.

In the model examining the mediator, sexual shape dimorphism, on the dependent variable, residualized attractiveness (Table 3 path b, Figure 4b), we controlled for RaterGen, as previous literature often suggests a preference for sex-typicality primarily concerning the opposite gender. The results, however, indicated that structural masculinity was generally associated with lower attractiveness (β_1j_ *= −*0.217 *[−0.233, −0.201]*, *t = −26.175*, *p < 0.001*, d = *−*0.27). This effect was somewhat stronger for female raters (Figure 4b), as the interaction term between RaterGen and Dimorphism was significant (β_3j_ *=* 0.071 [0.061, 0.081], *t = 13.303*, *p < 0.001*, d = 0.09). This aligns with several studies reporting similar outcomes (Perrett et al., 1998; Kočnar et al., 2019). A proposed explanation for this phenomenon is that for women, enhanced masculine facial characteristics can also lead to negative personal attributions, such as coldness, aggressiveness, or dishonesty, which are detrimental to romantic relationships and paternal investment. RaterGen itself had a small but significant impact on residualized attractiveness (β_2j_ *= −0.058 [−0.109, −0.006], t = −2.189*, *p = 0.028*, d = *−*0.07), a phenomenon we have previously identified as the GRG, with male raters generally providing more stringent ratings.

After accounting for sexual shape dimorphism, the direct effect of StimGen on residualized attractiveness (Table 3 path c’, Figure 4d) remained significant but was substantially reduced in magnitude (β_1j_ *= −*0.113 [*−*0.187, *−*0.039], *t = −*3.021, *p =* 0.002, d = *−*0.14), indicating partial mediation. To quantify the extent of mediation, we calculated the proportion of the total effect of StimGen on attractiveness that is explained by sexual shape dimorphism. Specifically, we computed the indirect effect (a × b), which quantifies how much of the relationship between the independent variable (StimGen) and the dependent variable (residualized attractiveness) is explained by the mediator (sexual shape dimorphism), and divided it by the total effect (c). Multiplying this quotient by 100 revealed that 66.2% of the total effect of StimGen on attractiveness was mediated by sexual shape dimorphism.

To assess the statistical significance of the indirect effect, we conducted nonparametric bootstrapping with 1,000 samples (Preacher & Hayes, 2008). This method resamples the dataset with replacement, refits the mediation models in each resample, and generates a distribution of the indirect effect, from which we extracted the 95% confidence interval. The results confirmed a significant indirect effect of StimGen on attractiveness through sexual shape dimorphism (*a × b = −0.25*), with a bootstrapped 95% CI of *[−0.264, −0.231]*, demonstrating that dimorphism significantly mediates the relationship between StimGen and attractiveness.

This model also controlled for RaterGen. The GRG (β_2j_ *= −0.177 [−0.236, −0.117], t = −5.816*, *p <0.001, d = −0.22*) and the interaction between StimGen and RaterGen (β_4j_ *= 0.204 [0.172, 0.237], t = 12.377, p <0.001, d = 0.26*) remained virtually unchanged compared to the total effect model (path c). Again, structural masculinity was generally associated with lower attractiveness (β_3j_ *= −0.298 [−0.322, −0.275], t = −25.067, p < 0.001, d = −0.38*). However, this time this effect did not vary by the raters’ gender (β_6j_ *= 0.001 [−0.018, 0.021], t = 0.147, p = 0.883*). Instead, it depended on StimGen, with increasing masculinity harming attractiveness more for female faces than male faces (β_5j_ *= 0.240 [0.205, 0.275], t = 13.613, p < 0.001, d = 0.3*). The three-way interaction between dimorphism, StimGen, and RaterGen was not significant (β_7j_ *= −0.009 [−0.041, 0.021], t = −0.612, p = 0.54*).

## Discussion

Our meta-analysis provides robust empirical evidence for a long-speculated phenomenon in the literature: female faces are perceived as more attractive than male faces, with an average rating advantage of 0.4 standard deviations. This GAP holds across rater genders, most cultural backgrounds, and ethnicities of portrayed individuals. Notably, our data suggest a possible deviation among African raters and African faces, though it remains unclear whether this reflects a genuine difference or is the result of sampling limitations.

Before exploring potential explanations for the GAP, we first consider whether methodological factors could have artificially produced this effect. One possibility is self-selection bias. In many studies on facial attractiveness, participants not only rate faces but also volunteer as models for the stimulus dataset, potentially introducing a gender-specific self-selection bias. If attractive women are more likely to participate in these photo sessions, while male participants do not exhibit a similar self-selection tendency, a spurious GAP could emerge. To evaluate this possibility, we examine the distribution of attractiveness scores in our harmonized dataset more closely (Figure 1). If self-selection bias were responsible for the GAP, we would expect an underrepresentation of low-attractiveness female faces, leading to a truncated left tail in the female distribution. Conversely, if unattractive men were disproportionately included in the dataset, we would expect an overrepresentation of low-attractiveness male faces, reflected in a local inflation in the left tail. However, both male and female distributions are symmetrically bell-shaped in their central range (Figure 1a,b), arguing against a self-selection bias. Rug plots along the x-axis suggest that male and female faces are similarly represented across the attractiveness spectrum.

Furthermore, Q-Q plots (Figure 1c,d) confirm that most attractiveness scores—particularly for female faces—closely follow a normal pattern. The only notable deviation occurs in the high-attractiveness range for male faces, where a localized inflation in the upper tail is observed (Figure 1d). This suggests that, if anything, highly attractive male faces may be somewhat overrepresented in the harmonized dataset (assuming that attractiveness is normally distributed in the population). Interestingly, despite the mild inflation in the upper tail, the male attractiveness distribution remains more concentrated around the mean (Figure 1b), with a sharper peak than the female distribution (Figure 1a). This suggests greater consensus among raters regarding male attractiveness, whereas female attractiveness ratings exhibit greater variability.

Together, these observations suggest that the processes of aesthetic judgment—rather than a sampling bias—is responsible for the GAP. Further supporting this conclusion, we observe the GAP consistently across individual study datasets, employing diverse stimulus selection procedures. This includes: self-selected participation (Fiala et al., 2021; Kočnar et al., 2019; Kleisner et al., 2024; Pavlovič et al., 2023; Qi & Ying, 2022; Torrance et al., 2014; Zhang et al., 2019), use of facial databases (Ebner et al., 2018; Chen et al., 2021; Germine et al., 2015; Hewer & Lewis, 2024; Leger et al., 2024; Lindeberg et al., 2019; Sutherland et al., 2017), author-selected images (Gouda-Vossos et al., 2016; Sutherland, Rhodes et al., 2020; Talamas, Mavor, & Perrett, 2016; Wassiliwizky et al., 2023), photographs from twin cohort studies (Mitchem et al., 2015), profile images from online platforms (Jaeger et al., 2019), stills from video files (Bowdring et al., 2021), and automated AI-mediated web searches (Sano & Kawabata, 2023). The replication of the GAP across such varied selection methods strongly suggests that the effect is genuine rather than a result of a sampling bias.

While self-selection is unlikely to explain the GAP, another potential artifact of data collection could be systematic differences in grooming practices between genders. Women typically invest more time and resources in grooming than men (Das & Stephen, 2011). However, most studies follow standardized photo-shooting protocols requiring participants to remove makeup and other enhancing accessories (e.g. Ebner et al., 2018; Kleisner et al., 2024; Torrance et al., 2014; Zhang et al., 2019). While some grooming practices, such as eyebrow shaping or long-term skincare routines, may not be entirely controlled by these protocols, their impact is likely minimal.

All of this naturally leads to the question: What explains the GAP? While evolutionary frameworks have traditionally been the dominant lens through which the GAP has been viewed— assuming its existence without direct empirical evidence—these theories focus exclusively on opposite-sex attraction, mate selection, and reproductive success. Within these theoretical boundaries, explaining the variation in same-sex ratings and the cultural differences in the GAP becomes challenging, suggesting that factors beyond biological predispositions also play a role. Given these limitations, sociocultural factors and norms merit further consideration. As noted earlier, female beauty is idealized in many cultures and reinforced by media, advertising, and societal expectations. Internalized beauty standards may foster unconscious biases, leading to, or amplifying, the observed difference. In other words, ratings of female images may blend subjective aesthetic responses to facial features with cultural expectations that link attractiveness more strongly with women. Disentangling these intertwined factors is crucial and requires targeted research to better understand the role of cultural expectations in shaping explicit attractiveness ratings. Furthermore, the cultural variations in the effect strength of the GAP observed in our ubiquity analysis (Figure 3) may be directly tied to how strongly beauty is associated with the female gender within a given culture—an area warranting further investigation.

A particularly intriguing aspect of the GAP is its interaction with the rater’s gender. Specifically, the difference in ratings between female and male faces is more pronounced among female raters. This moderation highlights fundamental differences in rating behavior between genders when evaluating facial attractiveness, particularly in same-sex and opposite-sex judgments. Notably, the unexpected rating behavior of female raters primarily drives the pattern observed in Figure 2a. While men, as anticipated, rate female faces more favorably than male faces, female raters exhibit the same preference—but to an even greater extent. As a result, male and female raters do not significantly differ in their ratings of male images. However, they diverge sharply in their ratings of female images, with female raters being notably more generous in evaluating their same-sex peers. This peer generosity persists even after adjusting for structural facial features (Figure 4d). These observations raise two key questions: Given that most raters in these studies identify as heterosexual, why does female rating behavior fail to align with their expected attraction to men? And why are women so generous in their evaluations of other women?

One contributing factor to women’s lower ratings for male faces may be rooted in the well-established, aforementioned evolutionary principle that women are generally more selective than men in mate choice. This greater choosiness may dampen the average attraction women feel toward potential mates, compared to men, who typically display less selectivity. Additionally, female raters may apply more complex evaluative criteria when assessing male faces, incorporating factors such as perceived trustworthiness or dominance (Sutherland, Burton et al., 2020), which could dilute purely aesthetic judgments. Social norms and self-presentation concerns may also play a role. Due to societal expectations that discourage overt displays of female desire, women may feel uncomfortable openly expressing attraction, particularly in settings where their responses are recorded and scrutinized. This could lead them to adopt a more reserved approach, resulting in more conservative ratings of opposite-sex faces. These factors are not mutually exclusive. They may interact and collectively contribute to the relatively lower ratings women assign to male faces.

In contrast, the overly generous ratings that women give to other women could stem from the cultural bias discussed earlier, which associates the female gender more strongly with beauty. This bias may be particularly pronounced among female raters, as studies have shown that across many societies, girls and women are socialized from an early age to place a high value on appearance (Calogero & Thompson, 2010; Hanson et al., 2024; Rice, 2014; Smolak & Murnen, 2011). Consequently, women are more frequently exposed than men to media that perpetuate idealized standards of female beauty. Additionally, given the societal pressures women face around appearance, this generosity might also reflect a sense of peer solidarity and a mutual recognition of shared pressures. Together, these factors could foster greater attunement to, and appreciation of, each other’s beauty, resulting in more favorable evaluations of female faces by female raters.

Importantly, the moderation of the GAP by the rater’s gender underscores that same-sex and opposite-sex ratings differ substantially in nature. In other words, while male ratings of attractiveness align with romantic, heterosexual attraction, this link is less clear for female raters. Technically, this challenges the principle of measurement invariance, as attractiveness ratings likely reflect different constructs depending on the rater’s gender. Such inconsistency complicates interpretive inferences and highlights the need for future research to address this issue. One potential solution could involve rephrasing the typical question from ‘How attractive is this face?’ to ‘How much do you feel physically attracted to this person?’

As a side note, our results do not support the assumption of intra-gender competition for either gender, which predicts that raters perceive same-sex peers as competitors and (subconsciously) downplay their attractiveness to enhance their own self-image or avoid feelings of inferiority. Instead, we observe the opposite for female raters, who evaluate other women very generously, and find no evidence that male raters tone down the attractiveness of their same-sex peers compared to female ratings of male faces (Figure 2a).

Our results reveal that men tend to evaluate facial attractiveness more stringently overall, although this GRG effect is relatively small compared to the GAP (d = 0.16). This tendency aligns with findings from other domains, where male raters have exhibited similar stringency in evaluations of life satisfaction (Plagnol & Easterlin, 2008; Zweig, 2015), perceptions of others’ personalities (Winquist et al., 1998; Srivastava et al., 2010), and online reviews (Bayerl et al., 2024). Among these, we focus particularly on the latter, online reviews, as they most closely resemble the aesthetic judgments analyzed in our study. The recent large-scale investigation by Bayerl and colleagues of gender-specific rating behavior across major platforms such as Amazon, IMDb, and TripAdvisor found that men consistently provided more stringent ratings than women for products, places, and services. Notably, the GRG effect for facial attractiveness observed in our study (d = 0.16) is larger than the effects reported there for products (d = 0.05) and travel destinations (d = 0.01) but similar to those reported for movies (d = 0.14) and music (d = 0.16).

To examine the consistency of the GRG across raters’ cultural backgrounds and the portrayed ethnicities, we conducted two ubiquity analyses similar to those for the GAP (reported in the Supplemental Material). The results showed that the GRG was far less consistent across cultural backgrounds and portrayed ethnicities compared to the GAP, with the effect even reversing direction in several cases (Figure S3). This also aligns with Bayerl et al.’s results, which have shown non-significant or reversed effects in certain countries and certain domains, such as no GRG effect for judging paintings or a reversed GRG in ratings for beauty and hair salons, where women were more stringent than men in their judgment. The variability across domains and contexts underscores the need for further research into the cultural and psychological mechanisms underlying this bias.

Our morphometric mediation analysis shows that two-thirds (66.2%) of the GAP between male and female faces can be attributed to differences in structural sex-typicality. Importantly, when comparing Figure 4c and Figure 4d, we observe that after controlling for sexual shape dimorphism, the attractiveness ratings of female images (orange bars) drop substantially, whereas male images (blue bars) remain virtually unchanged. This indicates that the mediation mechanism acts primarily on female faces, meaning that they lose a substantial portion of their advantage once structural dimorphism is accounted for. Furthermore, this suggests that the previous strength of the attractiveness gap was largely driven by a preference for structural femininity in female faces by both genders. In contrast, the lower attractiveness ratings of male faces were not strongly mediated by sexual shape dimorphism, indicating that their disadvantage is not attributable to structural sex-typicality. In fact, even after controlling for this factor, male faces still receive overall lower attractiveness ratings than female faces (Table 3 path c’). This is likely due to women still rating female faces significantly higher than male faces (’F : f’ vs. ‘M : f’ in Figure 4d), demonstrating a persistent peer generosity among female raters. Additionally, we observe that even after controlling for dimorphism, male raters remain much stricter when it comes to judging of female faces (’F : f’ vs. ‘F : m’), while no comparable male stringency bias is observed for male images (’M : f’ vs. ‘M : m’), suggesting that their evaluation criteria for male faces differ from those for female faces.

To conclude, the GAP is a robust finding with a substantial effect size, observed across diverse cultural and ethnic backgrounds. Its persistence, even after controlling for structural sex-typicality, suggests that the difference is not solely driven by facial structure but also influenced by the perceived gender of the face. The GAP’s dependence on the rater’s gender—driven by female raters’ peer generosity in same-sex evaluations and their lower-than-expected ratings for opposite-sex faces— adds complexity to the role of gender in attractiveness assessments. We propose that these biases stem from cultural norms linking femininity with beauty and a potential sense of solidarity among women. Recognizing gender-specific biases in attractiveness evaluations has significant academic value, informing research in fields such as evolutionary psychology, empirical aesthetics, sociology, cultural anthropology, and gender studies. Additionally, understanding these effects can enhance the design of future studies, improve rater selection, and refine the interpretation of results. Outside academia, our findings highlight how culturally embedded norms and gender-based expectations shape aesthetic judgments and influence societal perceptions. Finally, our results encourage further investigation into the cultural factors that influence aesthetic evaluations of human attractiveness.

## Acknowledgements

We would like to thank Brenda Edosomwan for her assistance with the literature search and Klaus Frieler for his valuable comments on our analyses. We are also deeply grateful to the corresponding authors who provided us with their original raw data.

## Author Contributions Statement

EW conceptualized the study, conducted the analyses, prepared the visualizations, and wrote the original draft. BPZ and FU reviewed and edited the manuscript. All authors contributed to the interpretation and discussion of the results and approved the final version for submission.

## Data and Code Availability Statement

All preprocessed data from our own studies are available on the Open Science Framework [link to be added]. The harmonized dataset can be reconstructed using the information provided in Table 1. [NB: We are seeking permission from all corresponding authors to make the full harmonized dataset available upon publication.] The R script for all statistical analyses, figures, and tables are also published on the Open Science Framework.

## Supplemental Material

**Fig.S1.**
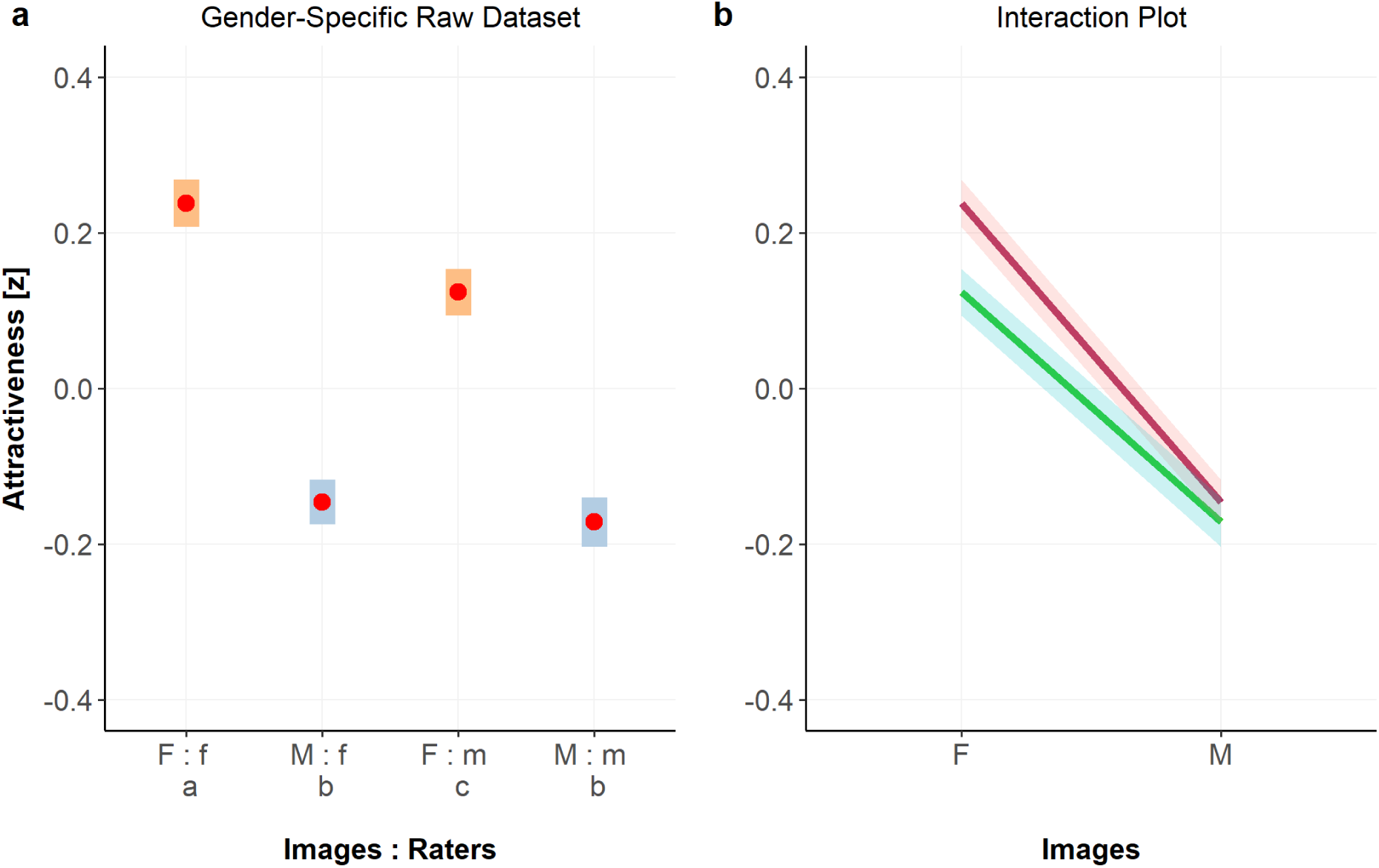
Meta-analysis results for the LMM of the harmonized raw dataset. Panel **a** shows estimated marginal means, including their 95% confidence intervals. The gender of the facial image is indicated in capital letters, while the gender of the rater is shown in lowercase letters. Conditions marked with different subscript letters are significantly different at a p-value of < 0.001. Panel **b** depicts the interaction effect between the image’s gender (on the x-axis) and the rater’s gender (line colors: maroon for female raters, green for male raters). Lines represent predicted attractiveness scores, the shaded ribbons indicate 95% confidence intervals. Note that the results pattern is almost identical to Fig.1a and 1c in the manuscript, demonstrating that the aggregation for the main analysis did not affect the results. For statistical details, please refer to Table S1.

**Table S1.** LMM results for the harmonized raw dataset. Attractiveness (z-scored) was predicted based on both the image’s gender (StimGen: M, F) and the rater’s gender (RaterGen: m, f). The following model with random intercepts for stimuli and individual raters was tested: Attractiveness_ij_ = β_i0_ + β_1j_ StimGen_ij_ + β_2j_ RaterGen_ij_ + β_3j_ (StimGen_ij_ x RaterGen_ij_) + ε_ij_.

| *Harmonized Raw Dataset* |  |  |  |
| --- | --- | --- | --- |
| *Predictors* | *Estimates*<br>*[95%-CI]*<br>*Westfall's d* | *t* | *p* |
| **Fixed Effects** |  |  |  |
| $\gamma_{00}$<br>(Intercept) | 0.238<br>[0.208, 0.268] | 15.408 | <0.001 |
| $\beta_{1j}$<br>StimGen:M | -0.384<br>[-0.413, -0.355]<br>d = -0.47 | -25.986 | <0.001 |
| $\beta_{2j}$<br>RaterGen:m | -0.114<br>[-0.144, -0.085]<br>d = -0.14 | -7.638 | <0.001 |
| $\beta_{3j}$<br>StimGen:M x<br>RaterGen:m | 0.089<br>[0.082, 0.096]<br>d = 0.11 | 25.098 | <0.001 |
| **Random Effects** |  |  |  |
| $\epsilon_{ij}$ | | 0.427 | |
| $V_{i0}$ | | 0.236<sub>Stim</sub> | |
| $V_{0j}$ | | 0.311<sub>Rater</sub> | |
| ICC |  | 0.562 |  |
| N |  | 5191<sub>Stim</sub> / 8986<sub>Rater</sub> |  |
| Observations |  | 830,204 |  |
| Conditional R<sup>2</sup> /<br>Marginal R<sup>2</sup> |  | 0.576 / 0.031 |  |

**Table S2.** Cross-tabulated frequency table. Number of stimuli by rater’s cultural region and stimulus ethnicity. Stimulus ethnicity could not be reconstructed for stimuli from four datasets (i.e., Studies 1, 6, 13, 20 and 21 in Table 1).

| **Rater’s Cultural Background** | **Stimulus Ethnicity** |  |  |  |  |  |
| --- | --- | --- | --- | --- | --- | --- |
|  | African | Asian | Caucasian | Latino | Middle-Eastern | Multi-Ethnic |
| Anglophone Countries | **300** | **412** | **3058** | **92** | **4** | **10** |
| Asia Eastern | 0 | **5059** | **1849** | 0 | 0 | 0 |
| Asia Southern | 0 | 0 | **120** | 0 | 0 | 0 |
| Europe Eastern | 0 | 0 | **794** | 0 | 0 | 0 |
| Europe Northern | 0 | 0 | **120** | 0 | 0 | 0 |
| Europe Southern | 0 | 0 | **120** | 0 | 0 | 0 |
| Europe Western | **22** | **36** | **2906** | **22** | **26** | **84** |
| Latin America | 0 | 0 | **120** | **138** | 0 | 0 |
| Middle East | 0 | 0 | **120** | 0 | **271** | 0 |
| Sub-Saharan Africa | **200** | 0 | **240** | 0 | 0 | 0 |

**Table S3.** Hierarchical model testing. The initial model (model.0) predicted Attractiveness by the factors StimGen and RaterGen. The consecutive models 1 to 3 append cumulatively one of the following covariates: sexual shape dimorphism, facial averageness, and youthfulness. The statistical model fit to the data is tested via the *anova* function in R. The best model performance is achieved when all three factors are included, highlighting their combined importance in explaining attractiveness.

| *Model Fit* |  |  |  |  |  |  |
| --- | --- | --- | --- | --- | --- | --- |
| *Model* | *n of parameters* | *added covariate* | *AIC* | *BIC* | *Chi squared* | *p* |
| model.0 | 7 | [StimGen * RaterGen] | 400657 | 400728 |  |  |
| model.1 | 8 | Dimorphism | 399847 | 399928 | 811.98 | <0.001 |
| model.2 | 9 | Averageness | 399113 | 399204 | 736.11 | <0.001 |
| model.3 | 10 | Youthfulness | 398692 | 398794 | 422.73 | <0.001 |

\* AIC: Akaike information criterion, BIC: Bayesian information criterion

**Figure S2.**
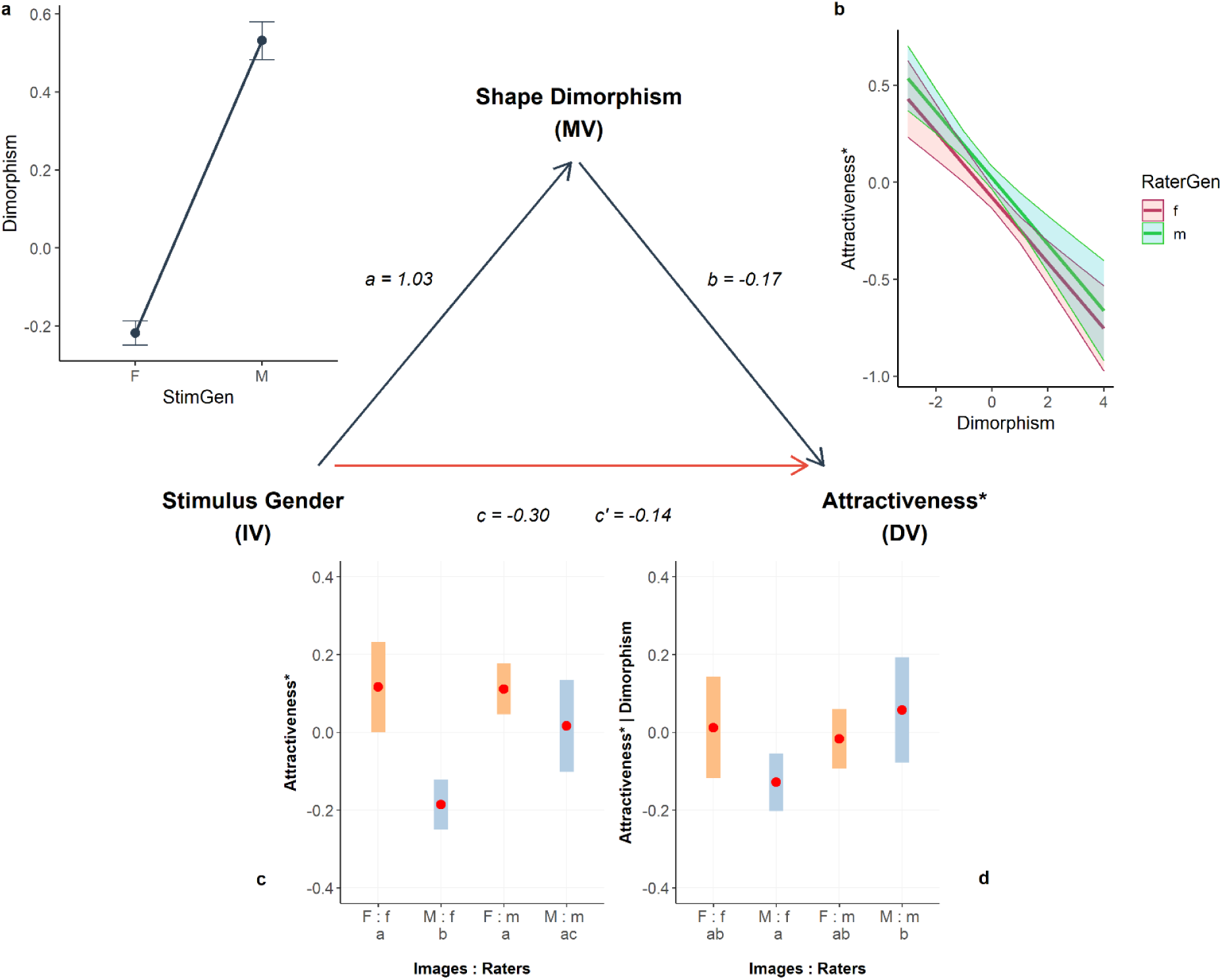
Replication of the multilevel mediation analysis using aggregated data. With StimGen as the independent variable (IV), Sexual Shape Dimorphism as the mediator (M), and residualized Attractiveness as the dependent variable (DV). Path coefficients (a, b, c, c’) are displayed along the respective arrows. Estimated effects plots (**a–d**) illustrate the modeled relationships. In panel **c**, conditions marked with different subscript letters are significantly different at *p* < 0.001; in panel **d**, at *p* < 0.05. The pattern of results remains consistent with Figure 4, showing partial mediation, with an indirect effect (a × b) of *−*0.173 (bootstrapped 95% CI = [*−*0.235, *−*0.138]). Sexual Shape Dimorphism mediates 57.72% of the total effect of StimGen on residualized Attractiveness. The considerably larger error bars in panels c and d, compared to those in the raw dataset in the manuscript, reflect the fact that fewer data points (i.e., two averages per stimulus) were available in this analysis.

## Ubiquity of the GRG

Using a similar statistical approach as we did with the GAP, we looked into the prevalence of the GRG across the raters’ cultural backgrounds and the ethnicities of the portrayed individuals. In other words, we asked: In which cultural regions were men stricter than women in their judgments of facial attractiveness? And, for which portrayed ethnicities did men tend to rate faces more stringently than women? (These questions could also be framed from women’s perspective.) To address these questions, we conducted two analyses: one testing whether the effect of RaterGen on attractiveness varied across cultural backgrounds of the raters and another testing whether the effect of RaterGen varied across portrayed ethnicities. As in previous models, stimuli were treated as Level 2 units and Bonferroni correction was applied to pairwise post-hoc tests to control for multiple comparisons.

The interaction between RaterGen and cultural region was significant (*F_19,9984_ = 5.093, p < 0.001, η_p_² = 0.007, f^2^ = 0.007*), indicating that the GRG varied across cultural contexts. Similarly, the second analysis revealed a significant interaction (*F_11,6396_ = 11.156, p < 0.001, η_p_² = 0.012, f^2^ = 0.012*), suggesting that the GRG also varied by the portrayed ethnicity of the depicted individual. Both effects were small.

Figure S3 reveals a mixed pattern of results, showing that the GRG varied not only in magnitude but also in direction across cultural regions and portrayed ethnicities. Specifically, male raters provided significantly more stringent attractiveness ratings in Sub-Saharan Africa and Anglophone countries, indicating a male stringency bias in these regions (Fig. S3a). Conversely, in the Middle East and Southern Europe, female raters were stricter than male raters, resulting in a female stringency bias. In the remaining regions, the contrasts were not significantly different from zero. A similar pattern emerged for portrayed ethnicities (Fig. S3b). Men rated African, Asian, and Caucasian faces more stringently than women, while they rated Latino and Middle Eastern faces more generously. For multi-ethnic faces, the contrast was not significantly different from zero.

**Figure S3.**
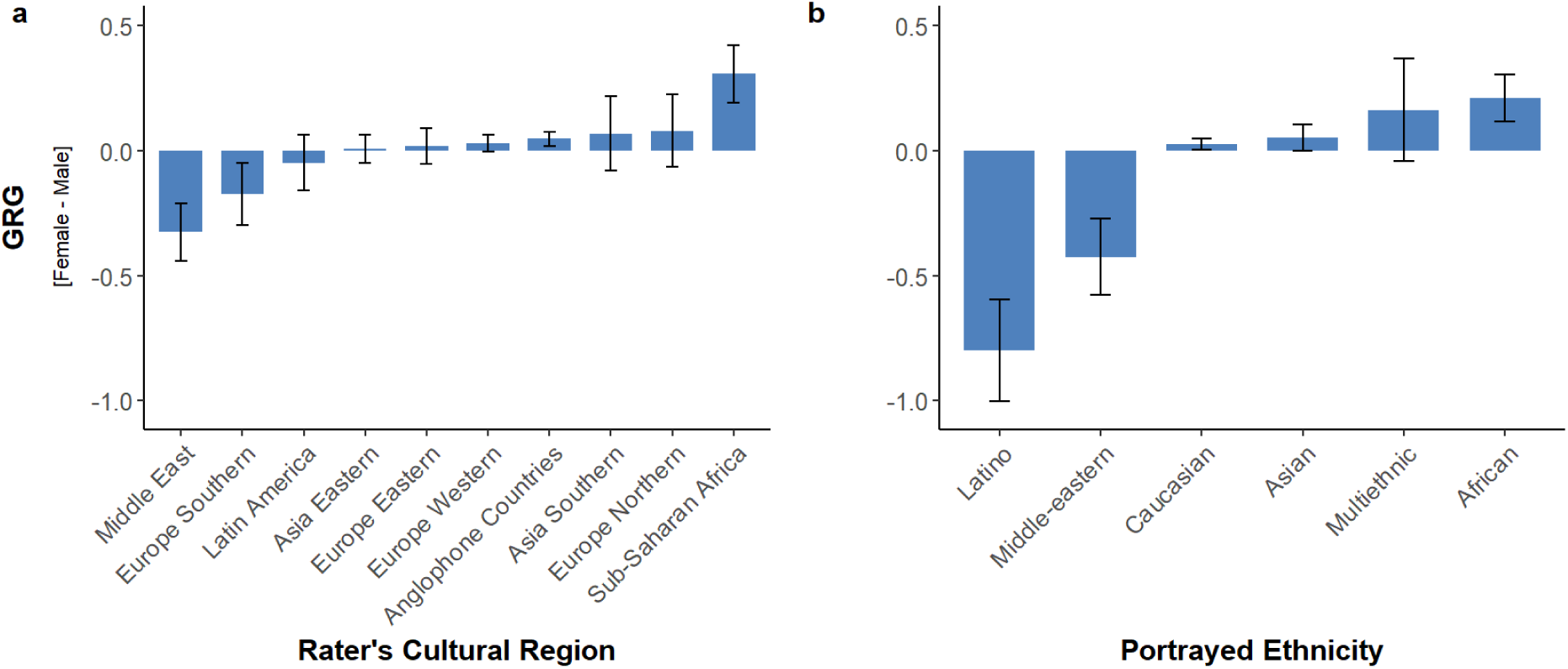
Ubiquity of the GRG. Panel **a**: Differences in attractiveness EMMs (z-standardized) across the cultural backgrounds of the raters, ordered by the magnitude and effect direction. The error bars reflect 95% confidence intervals. The GRG showed stricter ratings by male raters (at p < 0.001) in Sub-Saharan Africa and Anglophone Countries. In contrast, the GRG reversed direction in the Middle East and Southern Europe, where female raters were stricter (p < 0.001 and p < 0.01, respectively). For the remaining regions, differences were not significantly different from zero. Panel **b**: Differences in attractiveness ratings across portrayed ethnicities, ordered by the magnitude and effect direction. Stricter ratings by male raters were observed for African (p < 0.001), Asian (p < 0.05), and Caucasian (p < 0.05) faces. Conversely, the GRG reverses direction for Latino and Middle Eastern faces, where female raters were stricter (both p < 0.001).

